# Biomimetic Lipid-polymer composite membrane model to study Permeability through Cornea

**DOI:** 10.1101/2023.12.21.572515

**Authors:** Ruchira Chakraborty, Chavan Tejas Avinash, Manju Misra, Prasoon Kumar

**Affiliations:** BioDesign and Medical Devices Laboratory, Department of Medical Devices, National Institute of Technology Rourkela, Sundergarh -769008, Odisha, India; Department of Pharmaceutics, National Institute of Pharmaceutical Education and Research (NIPER)-Ahmedabad, Palaj, Gandhinagar-382355, Gujarat, India

**Keywords:** biomimetics, lipid-polymer composite, artificial cornea, permeability, drugs

## Abstract

The excised animal cornea is the gold standard for testing and evaluating drug permeability through a cornea. However, it has a concise shelf life and encounters ethical concerns. Also, ex-vivo models, which are incomplete replicas of the human cornea, may provide faulty results in the pre-clinical studies. To circumvent these problems, we have proposed an in-vitro biomimetic lipid-polymer composite membrane (BLCM) model as an artificial cornea to study drug permeability. We designed and fabricated a free-standing, electro-spun polystyrene (PS) nanofibrous membrane and impregnated its pores with phosphatidylcholine (PC). SEM, FTIR, and goniometer characterized the BLCM. Permeation data of the drug ganciclovir through the BLCM model in a Franz-diffusion cell corroborates with the excised goat corneal system. Also, owing to the simple and scalable fabrication method, BLCM can be used as an alternative to animal models for initial drug permeability screening and studies and accelerate drug development.

---

The human eye is a complex organ that comprises multiple physical and physiological barriers, reducing the bioavailability of drugs delivered through the eye. Different drug delivery routes have been explored for various diseases and discomforts of the eye (topical, intracameral, and subconjunctival). (Bachu et al., 2018) The topical route is the most popular owing to its non-invasiveness (painless route of drug administration), high patient acceptability, and the possibility of self-administration of the drugs. (Nayak and Misra, 2018) The correcst permeability prediction for the drug candidates and ophthalmic formulations is a crucial step in the drug development process, mostly done with the help of excised animal corneas of rabbit, pig, goat, or bovine sources. (Agarwal and Rupenthal, 2016) Though highly accurate, these models are costlier and have a shorter shelf life. Further, animal ethics-related issues and more complex permeation mechanisms discourage their usage. Also, incomplete replicas of the human cornea may sometimes provide faulty results in pre-clinical studies.(Subasinghe et al., 2021) On the other hand, cell monolayers and tissue-engineered models provide a range of applications, but they sometimes prove to be cumbersome and tricky to handle. (Bittermann and Goss, 2017) Further, considering the limited availability of human-derived primary cells, these approaches are not optimal for permeability studies. (Agarwal and Rupenthal, 2016; Reichl et al., 2004)

Conversely, cell-free models are composed primarily of phospholipids that mimic the cell membrane of the epithelial cell layer since they are the primary barrier to drug permeation.(Berben et al., 2018) These are simple models efficient for drug permeability studies. Permeapad and PAMPA are examples of cell-free models that have proven successful in providing high throughput screening of drugs across the membrane. The Permeapad model comprises a phospholipid layer sandwiched between two support layers composed of cellulose.(Jacobsen et al., 2020) The PAMPA membrane comprises phospholipids dispersed in organic solvents such as dodecane and suspended in a support layer. It overcomes the complexities of active transport and enables the kinetics of test compounds on simple permeation properties. (Avdeef, 2005) Given that the PAMPA membranes mimic the in-vivo condition of the epithelial interphases, it is vital to withstand body fluids such as bile, enzymes, FaSSIF, or FaHIF. However, in the previous PAMPA models, environmental conditions such as pH, excipients, and biomimetic media affect functional stability and permeation, reducing functionality. (Berben et al., 2018) Additionally, it has reduced stability at temperatures higher than 37°C and cannot withstand mechanical stresses, whereby increased leakage can be observed. (Berben et al., 2018) Permeapad overcomes these structural shortcomings with high robustness and decreased temperature sensitivity but is costly and limited for hydrophilic drugs due to low solubility.(Jacobsen et al., 2023) This necessitates a mechanically stable, accurate model predicting drug permeability through the corneal epithelium.

In this model, we have optimized the different concentrations of phosphatidylcholine impregnation in a polystyrene nanofibrous membrane matrix to develop a biomimetic lipid-polymer composite membrane (BLCM) to mimic the corneal barrier layers (Figure 1A). The BLCM model described here is a simple yet scalable, thermostable, easy-to-handle model for emulating a corneal epithelial barrier. We have characterized BLCM and demonstrated that the flux of a hydrophilic ganciclovir drug through the BLCM membrane corroborates with the permeability of the drug obtained from the excised goat corneal system. The distribution and arrangement of amphiphilic phospholipid molecules in the hydrophobic polystyrene bed provide a new understanding of the permeation mechanism, making the proposed membrane a possible alternative for initial drug permeability studies. Polystyrene (PS) (Himedia, CAS No-9003-53-6), tetrahydrofuran (THF) (Himedia, CAS No-AS-071), N, N Dimethylformamide (DMF) (Sisco Research Laboratories, 68-12-2), cellulose acetate sheet (CS), phosphatidylcholine (PC) (Sisco Research Laboratories, 8002-43-5) were used for the research.

**Figure 1:**
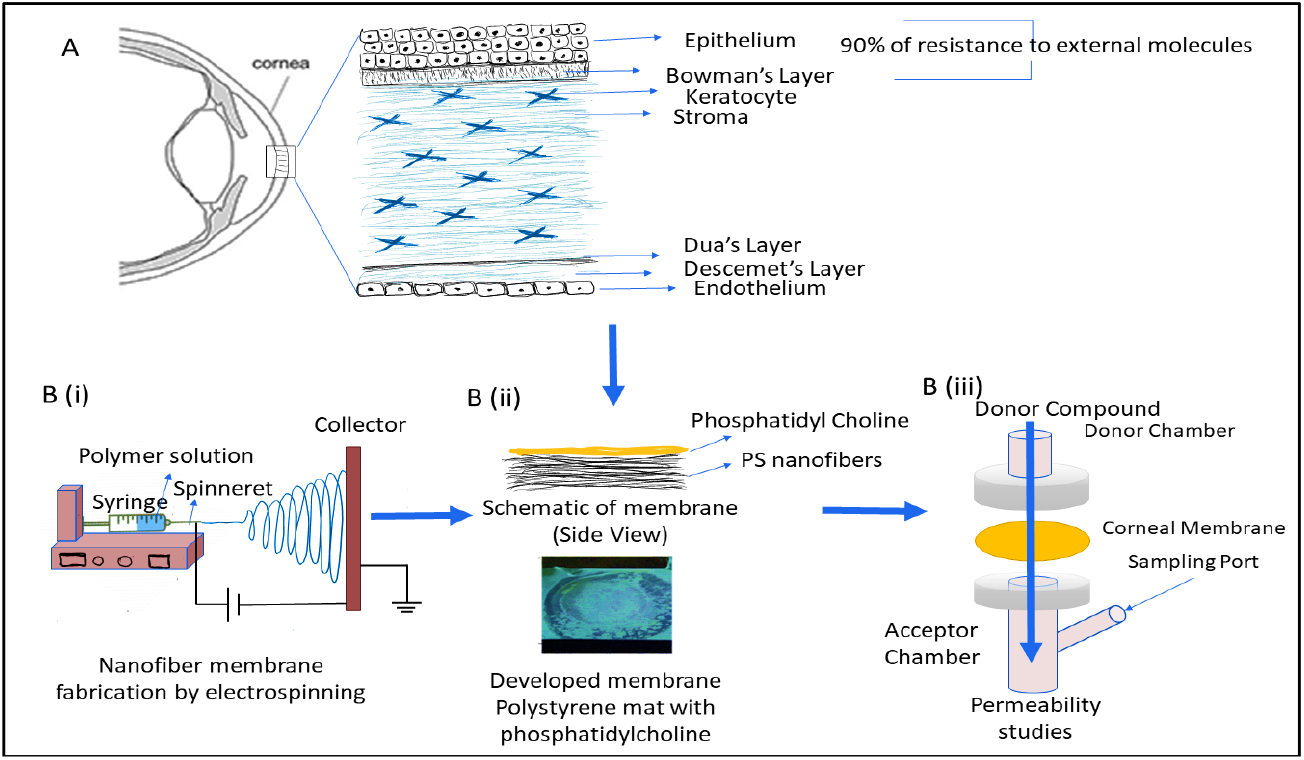
A schematic of the fabrication process (A) Different layers of the human cornea. (B(i)) Electro-spun nanofibrous polystyrene (PS) membrane fabricated. (B(ii)) Phosphatidyl Choline (PC) was poured over the random nanofibers (B(iii)). Permeation of drug molecules through the corneal membrane fabricated with PC entrapped in PS nanofibrous membrane.

A 15% wt./vol polystyrene solution was prepared in the THF and DMF solvent system (3:1). The solution was stirred over a magnetic stirrer for 6-8 hours before electrospinning. The homogenous, transparent polystyrene solution was loaded in a 5ml syringe and electrospun for 2.5-3 hours to generate a free-standing nanofibrous matrix. The optimized parameters for electrospinning include 15KV voltage between electrodes, 0.5ml/hr flow rate, 15 cm distance between the needle and collector electrode, and 24G needle (Nitanan et al., 2012). The substrate used was the circularly punched cellulose acetate sheet. The nanofibers obtained were examined for their morphological analysis through SEM (Figure 1B(i)).

The different weights of PC (5mg, 10mg, and 20mg) were mixed in 150ul of methanol. After that, the solution was poured onto the free-standing Electrospun nanofibrous PS membrane collected on a cellulose acetate paper and allowed to dry at an ambient condition for 12 hours to remove the residual solvent (Figure 1(B(ii))). The final PC content in the membrane was determined by taking the membrane weight before and after the entrapment of PC. (Pulugu et al., 2022) These dried membranes were then used for the permeation studies of the model drug. Samples were initially sputter coated with gold to obtain SEM images and were imaged at the accelerating voltage of 5 KV. The film was observed under a scanning electron microscope (SEM) (Carl Zeiss, Germany) to see the uniform distribution of PC into PS. SEM images were taken before and after PC entrapment and permeation studies. The transverse images of the membrane were also taken to determine its thickness. The contact angle was measured to confirm the PC entrapment into PS and mimic the cornea. Contact angle measurement was performed using the instrument KRÜSS drop shape analyzer. A sessile drop of water was used to measure the contact angle. A 3 μL water drop was placed on the sample using a syringe installed with a needle. The contact angle was measured directly by the software KRÜSS advanced 1.5.1.0. Attenuated total reflectance (ATR) spectroscopy was performed to understand the interaction between electro-spun PS fibers and PC. ATR of electro-spun PS fibers, PC, and PC-loaded PS was done to detect the functional groups’ presence and evaluate the molecular interactions. The examination was done from the range of 4000-400 cm^-1^ and was compared with the literature. (Cagnasso et al., 2010; Uche et al., 2020)

Permeation studies of the model drug (GCV) were performed using the proposed membrane and ex-vivo model. The GCV drug solution was prepared in PBS at 1mg/ml concentration. Permeation studies were performed in triplicate using the selected model and ex-vivo cornea using Franz diffusion cell assembly. For *Ex-vivo* studies, the whole eyeball of a goat was procured from the local slaughterhouse and was used within 1 hour of sacrificing the animal. The eye was kept in PBS buffer at pH 7.4 and 4°C. For the study, the cornea was separated from the eyeball and was cleaned thoroughly to remove any other adhered tissue using a standard solution. (Bhosale et al., 2022) The goat cornea was uniformly spread between the acceptor and donor compartment, as shown in Figure 1(B(iii)). After that, 14ml of PBS buffer at pH 7.4 was placed into the receptor compartment such that the cornea was in close contact with the buffer media. Thereafter, 2 ml of GCV solution (1 mg/ml) was placed in the donor compartment, and then the diffusion assembly was placed into an orbital shaker with continuous stirring at 30 rpm. The temperature of the diffusion assembly was maintained at 37°C. After that, fresh aliquots were taken from the receptor compartment at defined intervals and replaced with a fresh medium of PBS pH 7.4. The aliquot samples were analyzed using a UV-Visible spectrophotometer at 254nm. (Al-Badr and Ajarim, 2018) Similar studies for GCV solution were also performed using a biomimetic lipid-polymer composite membrane (BLCM) prepared using 10 mg of PC. The steady-state flux and apparent permeability coefficient were determined to compare the permeation through the membrane and *ex-vivo* corneal membrane.

The polystyrene (PS) nanofibrous matrix has nanofibers with smooth morphology and completely random orientation, as shown in Figure 2(A) The nanofibers deposited on a circularly punched cellulose acetate sheet created a free-standing nanofibrous membrane. The PC solution in methanol could reach and fill the pores of the free-standing nanofibrous membrane (Figure 2 (A, B)). The SEM images confirmed that PC was uniformly distributed into the electro-spun matrix. The thickness of the membrane was measured by taking the transverse SEM image (Figure 2(B(ii))) of the membrane. The thickness of the membrane was found to be in the range of 45μm and 75μm (Figure 2B(ii)). The thickness of corneal epithelium (CE) is also around 60μm. (Reinstein et al., 2008) The SEM image of the membrane was also taken after the completion of permeation studies and the drying of the membrane (Figure 2(C)). The image showed pores in the membrane, which might be due to the amphiphilic nature of PC since the study was carried on for an extended period, i.e., for 8 hours, compared to the regularly used study duration of 4 hours. The contact angle of the membrane before and after the PC entrapment. The contact angle measurement on the PS nanofiber membrane was 127.97±1.67°, suggesting it is highly hydrophobic. It was observed that after PC entrapment, the contact angle drops to 24.47±1.36^o,^ rendering it hydrophilic as observed in a native cornea (55-65°) Figure 2 (D, E). (Cope et al., 2009) This is only possible if PC molecules interact with the PS membrane such that hydrophilic moiety on PC is exposed to the outer surface. At the same time, the hydrophobic groups are buried inside the membrane’s pores. We propose the configuration of the molecules, as shown in Figure 2(E(iii)), to account for the hydrophilic and hydrophobic behavior of the BLC membrane. ATR was performed to determine the interaction between PC and PS. ATR of the PS matrix, PC, and PC-loaded PS matrix is shown in Figure 2(F), and it is evident from the ATR spectra that all the significant peaks of PS and PC have been retained in the PC-loaded PS matrix, indicating the absence of interaction between the PC and PS. This indicates the physical entrapment of PC in PS membrane.

**Figure 2:**
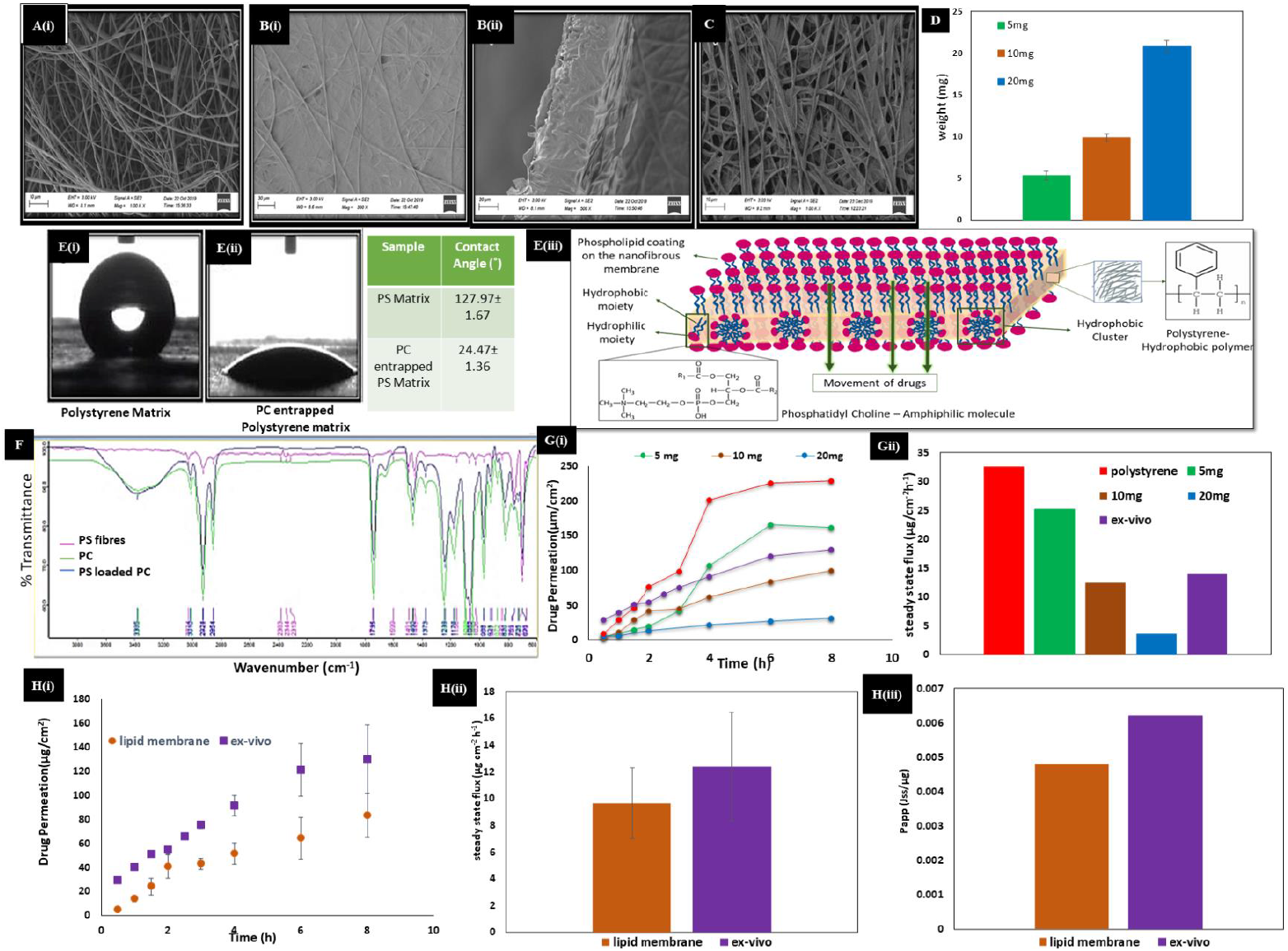
Electro-spun polystyrene matrix: A(i) Before Lipid entrapment, B(i) After lipid entrapment (surface view) and B(ii) After lipid entrapment (Transverse view), (C)After permeation studies (D) Lipid content of film (E) Contact Angle Measurement (E(iii)) Schematic Explanation of the dual nature of the membrane seen in contact angle measurement (F)ATR to study the interaction between Lipid and Polystyrene (G)(i) Comparative permeation studies from membrane containing different amount of PC, pristine PS and ex-vivo corneal membrane G(ii) Steady-state flux of GCV solution from membranes containing different amount of PC, pristine PS and ex-vivo corneal membrane (H(i)) Permeation Curve of GCV from membrane containing 10 mg PC, ex-vivo corneal membrane (H(ii)) Comparison by Steady state flux of GCV from membrane containing 10mg PC and ex-vivo corneal membrane n=3 (H(iii)) Comparison by Apparent permeability coefficient of GCV from membrane containing 10mg PC and ex-vivo corneal membrane

To optimize a membrane mimicking CE, we used a model drug, ganciclovir (GCV), a highly water-soluble drug (log P value of -1.66). (Heinig et al., 2011) It treats ocular diseases such as cytomegalovirus retinitis. (Jabs et al., 1987) The rate-limiting barrier for its ocular permeation is the Corneal Epithelium. Hence, the permeation of GCV through the BLCM model with varying PC content was compared with the ex-vivo corneal membrane to arrive at the optimal composition of the BLCM. It was found that the steady-state flux of GCV demonstrates that the pristine PS showed much higher permeation than that of the *ex-vivo* cornea. However, among the films of 5mg, 10mg, and 20mg PC, the film containing 10mg PC mimics the ex-vivo corneal permeation more closely (Figure 2 (G(i, ii))). We experimented with the BLCM model with 10mg of PC in the PS nanofibrous membrane. The measurement of drug concentration with time in the receiver compartment with the BLCM model and ex-vivo goat cornea as shown in Figure 2(H(i)) showed good similarity. The apparent permeability coefficient and steady-state flux were determined to quantify the trend further. It was shown that PC film containing 10mg of PC showed steady-state flux and apparent permeability coefficient values 12.39 μg cm^-2^h^-1^ and 0.0062 Jss/μg, respectively, which were statistically non-significant (p-value >0.05) to that of values obtained from ex-vivo studies 9.53 μg/cm^-2^h^-1^ and 0.0048 Jss/μg (Figure 2 (H(ii, iii). The optimization studies of GCV indicated that the model could be used to predict the permeability of hydrophilic drugs. The model needs to be validated for permeability using different classes of drugs with different ADMET properties. The mechanism of drug permeation through the BLCM model needs to be investigated in detail in future studies.

The alternative approaches to animal models for testing drugs during drug discovery have been researched for a long time. However, advanced materials have paved the way for devices, tissue-engineered models, and material platforms as alternatives to animal models. The electrospun PS membrane impregnated with PC led to the development of a barrier membrane (BLCM) mimicking the permeability properties of native goat cornea. We optimized the PC loading into the PS matrix and observed that 10% PC mimics goat CE. The permeability behavior of the proposed BLCM was tested and validated using a model drug, ganciclovir, and compared with the native goat cornea. The organization and distribution of phospholipids in the hydrophobic polystyrene membrane is responsible for the increased solubility of the hydrophilic drug in the membrane, allowing its permeation, just as in the case of in-vivo cornea. Further, the model must be optimized and tested for various drugs with different ADMET (absorption, diffusion, metabolism, excretion, and toxicity) properties. Also, understanding the permeation dynamics can be enriched with different drugs with different surface properties.

## Acknowledgment

TAC and MM would like to acknowledge NIPER-A for providing financial support for this project. RC would like to thank NIT Rourkela for her fellowship support. PK would like to thank NIPER-A, NIT Rourkela, and SRG Grant(SRG/2021/000859) by SERB, DST, Govt. of India.

